# Limited Introgression Supports Division of Giraffe into Four Species

**DOI:** 10.1101/303057

**Authors:** Sven Winter, Julian Fennessy, Axel Janke

**Author notes:** corresponding author, Contact:, Sven Winter*, Julian Fennessy, Axel Janke.

## Abstract

All giraffe (*Giraffa*) were previously assigned to a single species (*G. Camelopardalis*) and nine subspecies. However, multi-locus analyses of all subspecies have shown that there are four genetically distinct clades and suggest four giraffe species. This conclusion might not be fully accepted due to limited data and lack of explicit gene flow analyses. Here we present an extended study based on 21 independent nuclear loci from 137 individuals. Explicit gene flow analyses identify less than one migrant per generation, including between the closely related northern and reticulated giraffe. Thus, gene flow analyses and population genetics of the extended dataset confirm four genetically distinct giraffe clades and support four independent giraffe species. The new findings call for a revision of the IUCN classification of giraffe taxonomy. Three of the four species are threatened with extinction, mostly occurring in politically unstable regions, and as such, require the highest conservation support possible.

## Introduction

Traditionally, giraffe were classified as a single species (*Giraffa Camelopardalis*) with up to eleven subspecies proposed (Lydekker, 1904). However until recently, the classification into nine subspecies was most widely accepted (Dagg & Foster, 1976). It has been shown that in captivity some giraffe subspecies hybridize (Gray, 1972; Lackey, 2011; Lönnig, 2011), which seemed to supported the traditional single species concept for giraffe. However, multi-locus analyses of wild giraffe nuclear loci identified four monophyletic, distinct and evolutionary old groups of giraffe that should be recognized as four distinct species (Fennessy et al., 2016). This finding conflicts with former classifications and has been questioned based on the limited interpretation of traditional data e.g. pelage pattern, number of ossicones and geographic distribution (Bercovitch et al., 2017). The initial findings of four giraffe species (Fennessy et al., 2016) could better be criticized, because it did not involve explicit gene flow analyses. Imperative to understanding speciation in general from a genetic perspective is gene flow analyses, especially as the keystone of the biological species concept (BSC) is reproductive isolation (Coyne & Orr, 2004). The BSC implies that there is no or only very limited gene flow between species. It has been proposed that one or a limited number of effective migrants (up to 10) per generation (N_e_m) avoids genetic differentiation of populations and escapes a substantial loss of genetic diversity for neutral traits (Lacy, 1987; Mills & Allendorf, 1996; Vucetich & Waite, 2000; Wright, 1969). Thus, it is a conservative estimate that limited gene flow of less than one migrant per generation (N_e_m < 1) can lead to speciation, despite the occurrence of hybridization between mammal species. As shown, the BSC might need to be revised as some species naturally hybridize in the wild and produce fertile offspring e.g. bears (Arnold, 2016; Kelly, Whiteley, & Tallmon, 2010; Kumar et al., 2017) and whales (Bérubé & Aguilar, 1998; Spilliaert et al., 1991), and divergence can occur under genetic exchange (Arnold, 2016). Whilst the distinction of four giraffe species is consistent with population genetic analyses (Fennessy et al., 2016), gene flow among giraffe species has not yet been sufficiently analyzed. Here, we revisit the hypotheses of four giraffe species using population genetic methods that explicitly involve gene flow analyses with an increased dataset of 21 nuclear loci and 137 giraffe individuals from 21 locations across Africa (Fig. 1, Supplementary Table 1).

**Fig. 1.**
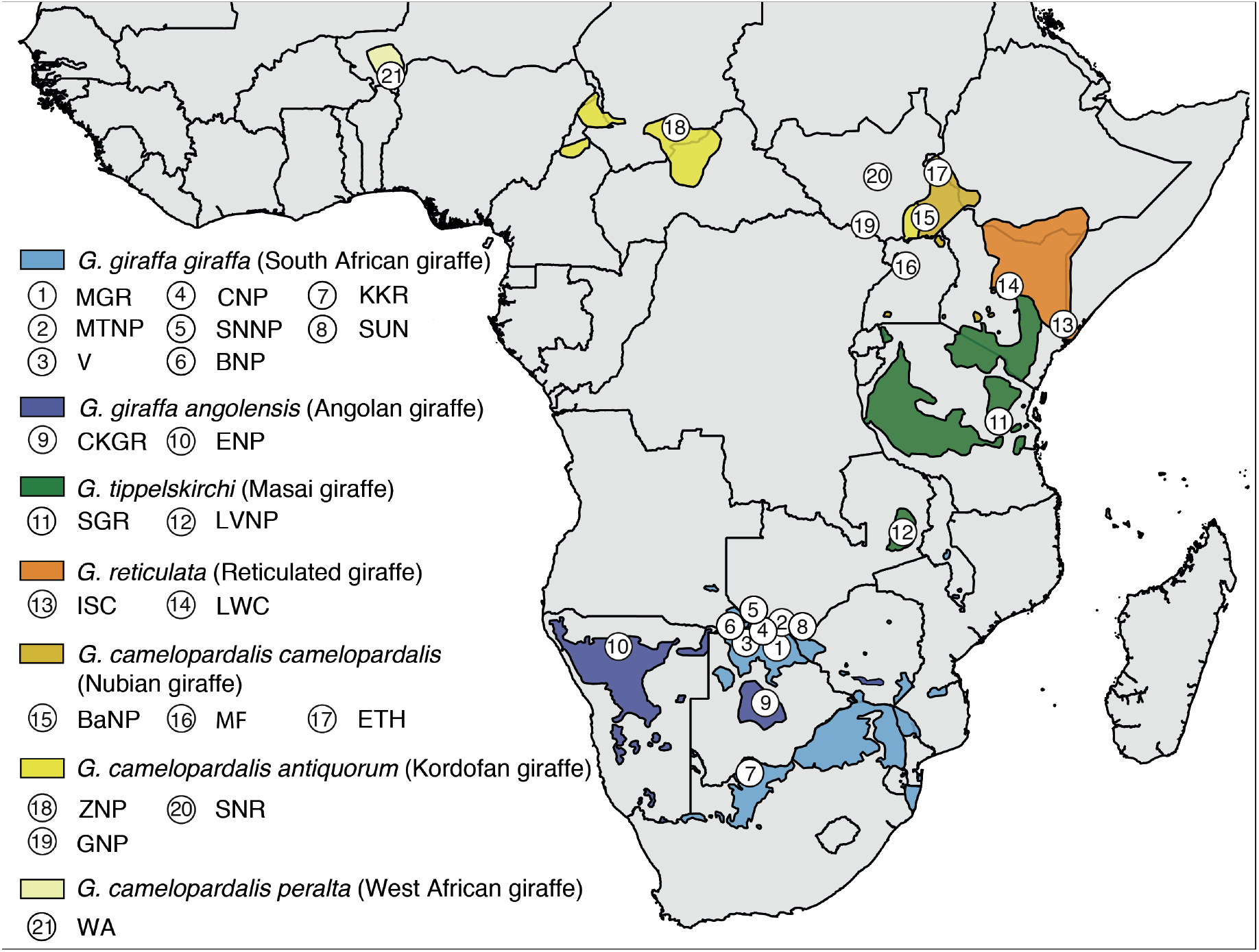
Map of Sub-Saharan Africa with giraffe (sub)species distributions and sampling locations. Geographic ranges (colored shadings) of giraffe as identified by the Giraffe Conservation Foundation (2017) were plotted on a map of Sub-Saharan Africa. Numbered circles represent sampling locations (for details see Supplementary Table 1). Species and common names as per Fennessy et al. (2016).

## Materials and Methods

### Sampling and DNA Extraction

Tissue samples from all giraffe species and five subspecies were collected by the Giraffe Conservation Foundation (GCF) and partners using remote biopsy darts with country-specific research permits between 2009 and 2016 in accordance with ethical guidelines and regulations of the respective governments and institutions. All samples were stored either in RNAlater (Invitrogen) or > 95% ethanol. Additional giraffe samples were added to the dataset of Fennessy et al. (2016) resulting in a total number of 217 giraffe individuals, mostly southern giraffe. The geographical origins and individual IDs are shown in Supplementary Table 1. Sample locations and geographical distributions are shown in Fig. 1. Additional southern giraffe individuals were only included if they are from a hitherto unrepresented region. DNA was extracted using either a Macherey-Nagel NucleoSpin Tissue Kit or a standard phenol-chloroform extraction method. All experimental protocols are in compliance with the guidelines for the best ethical and experimental practices of the Senckenberg Society, as well as with national guidelines of the respective countries.

### Amplification and sequencing

We PCR amplified and sequenced the seven intron markers previously published (Fennessy et al., 2016) for 32 new individuals and developed 14 additional intron markers as described (Fennessy et al., 2016). The 14 new intron markers were amplified and sequenced for a total number of 137 individual giraffe and the okapi (*Okapia johnstoni*). PCRs were performed with 10 ng genomic DNA giraffe and okapi specific primers (see Supplementary Table 2 for primer sequences and PCR conditions). We also amplified and sequenced the mitochondrial cytochrome b and control region for all new individuals as described previously (Bock et al., 2014). Each PCR was examined using agarose gel electrophoresis on a 1% agarose gel with ethidium bromide.

Sanger sequencing was performed for the forward and reverse strand using the BigDye terminator sequencing kit 3.1 (Applied Biosystems) with 5 ng of PCR product for each reaction and analyzed on an ABI 3730 DNA Analyzer.

The sequences were manually edited and aligned in Geneious v.6.1.8 (Kearse et al., 2012). Heterozygous insertions/deletions of nuclear sequences were resolved by hand or using Indelligent v.1.2 (Dmitriev & Rakitov, 2008) and verified by allele-specific primers if necessary. PHASE implemented in DnaSP v.5.10.01 (Librado & Rozas, 2009) was used to derive the allele haplotypes of the nuclear sequences using a threshold of 0.6 and allowing for recombination. All analyses, except of a mtDNA tree analysis, were performed using only nuclear allele haplotype data.

### Tree Analyses

The mitochondrial cytochrome b and control region sequences of 217 giraffe including newly generated as well as already published sequences (Bock et al., 2014; Brown et al., 2007; Fennessy et al., 2016, 2013; Hassanin et al., 2012, 2007; Winter, Fennessy, Fennessy, & Janke, 2018) (Supplementary Table 1) were aligned, as well as concatenated, and a Neighbor-Joining analysis was reconstructed in Geneious v.6.1.8 (Kearse et al., 2012). We used the HKY model of sequence evolution (Hasegawa, Kishino, & Yano, 1985), as suggested by jModelTest v.2.1.1 (Darriba, Taboada, Doallo, & Posada, 2012) with 1,000 Bootstrap replicates and sequences of two okapis were used as an outgroup.

A multi-locus Bayesian phylogenetic tree of the 21 intron markers for 137 individuals and the okapi as outgroup was generated with the StarBEAST2 (Ogilvie, Bouckaert, & Drummond, 2017) package in BEAST v.2.4.5. (Bouckaert et al., 2014, p. 2) under the JC model of nucleotide evolution as suggested as best fitting model by jModelTest v.2.1.1 (Darriba et al., 2012). A lognormal relaxed clock was used with 10^9^ generations and sampling every 20,000^th^ iteration. Convergence of the MCMC runs was analyzed with Tracer v.1.6.0 (Rambaut, Suchard, Xie, & Drummond, 2014), and TreeAnnotator v.2.4.5 (Rambaut & Drummond, 2016) was used to construct a maximum clade credibility tree with 30% burn-in for the nuclear markers.

### Population Genetic Analyses

Haplotype information for each locus deduced by DnaSP (Librado & Rozas, 2009) was used to code each individual. The haplotype matrix was then used to infer admixture with the Bayesian clustering algorithm implemented in STRUCTURE v.2.3.4. For the maximum number of populations (K) between 1-10, we sampled 250,000 steps following a 100,000-step burn-in, with 40 replicates each. The CLUMPAK webserver (Kopelman, Mayzel, Jakobsson, Rosenberg, & Mayrose, 2015) was used to average the results, and to infer the most likely K based on the posterior probability of K (Pritchard et al., 2000) and ΔK (Evanno et al., 2005). Additionally, the most likely K was deduced based on the estimated Ln probability of data (Ln Pr(X|K)) (Pritchard et al., 2010) using Structure Harvester (Earl & vonHoldt, 2012). Principal Component Analyses were performed with the R package adegenet (Jombart, 2008) in R v.3.2.3 (R Core Team, 2015) to assess the degree of similarity between defined population scenarios. Pairwise Fixation index (Fst) values were calculated in Arlequin v.3.5.2.1 (Excoffier & Lischer, 2010) based on the nuclear haplotypes.

### Gene flow analyses

Long-term average gene flow among and within the giraffe species were calculated in the coalescent genealogy sampler MIGRATE-N v.3.6.11 (Beerli, 2006; Beerli & Felsenstein, 2001) by estimating the mutation-scaled population sizes (Θ) for each population and migration rates (M) for each direction between a pair of populations. We used the Brownian motion mutation model and the Bayesian inference analysis strategy, as some parameter combinations are better estimated using the Bayesian approach compared to the Maximum-likelihood approach (Beerli, 2012). The transition/transversion ratio was set to 2.31 as estimated in MEGA v.7.0.16 (Kumar, Stecher, & Tamura, 2016) based on a concatenated alignment of all 21 loci. Variable mutation rates were considered amongst loci. We used the default settings for the Θ uniformpriors, and adjusted the M uniformpriors (0; 5,000; 10,000; 1,000), because the upper prior boundary appeared to be too small in initial first analyses. Several short-runs were performed to check for convergence of the runs. A long-chain run was performed for 6 million Markov Chain Monte Carlo (MCMC) iterations (60,000 recorded steps) and a burn-in of 600,000 iterations. An adaptive heating scheme was used with four chains and temperatures set by default with a swapping interval of 1. Convergence of the runs was checked by the posterior distributions, Effective Sample Size (ESS) and consistency of results between runs. In addition, we estimated short-term gene flow, as well as the probability of recent hybridization for each individual in BayesAss v.3.0.4 (Wilson & Rannala, 2003) using 100 million MCMC iterations, a burn-in of 10 million and a sampling interval of 1000 iterations. Mixing of the chain was improved by adjusting the acceptance rates for proposed changes to the parameters (allele frequencies and inbreeding coefficient) by adapting the mixing parameters for allele frequencies (ΔA) and inbreeding coefficients (ΔF) to 0.30. Convergence was checked in Tracer v.1.6.0 (Rambaut et al., 2014) and by consistency of results of several runs with different initial seeds. Results for short-term gene flow were visualized in circos plots using the Circos Table Viewer v.0.63-9 (Krzywinski et al., 2009).

### Calculation of gene flow rate

We calculated the effective number of migrants per generation (N_e_m) or rate of gene flow using two different methods. The first method to calculate N_e_m was based on the pairwise F_st_ values using equation (1) by Wright (1951):

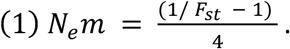

In addition, we calculated the N_e_m using the coalescent-based estimates for the mutation-scaled population size Θ and the mutation-scaled immigration rate M derived from MIGRATE-N. For autosomal markers equation (2) expresses the relationship between Θ_j_ (population size of the population receiving migrants) and M_ij_ (corresponding migration rate) (Marko & Hart, 2011):

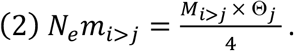

## Results

The putatively independent 21 nuclear gene loci are on different chromosomes or are clearly separated from each other in the bovine genome, a close relative with available chromosome level genome data (Supplementary Table 2). A Bayesian multi-locus tree analysis of the 21 nuclear loci (total of 16,969 nucleotides) for 137 giraffe, including all traditionally recognized giraffe subspecies (Fig. 2a), implies a clear separation into four giraffe clades: (1) a northern giraffe cluster including West African (*G. c. peralta*), Kordofan (*G. c. antiquorum*), and Nubian giraffe (*G. c. Camelopardalis*) which includes the former Rothschild´s giraffe (*G. c. rothschildi*), (3) the reticulated giraffe (*G. reticulata*), (3) the Masai giraffe (*G. tippelskirchi*) including former Thornicroft´s giraffe (*G. c. thornicrofti*), and (4) a southern giraffe cluster (*G. giraffa*) including South African (*G. g. giraffa*) and Angolan giraffe (*G. g. angolensis*). The monophyly of each of these four clades is supported by a posterior probability of p ≥ 0.95. However, in the analyses the exact relationships of southern and Masai giraffe relative to northern and reticulated giraffe could not be determined with significant probability (p ≈ 0.81) (not shown). A mtDNA Neighbor-Joining tree (Supplementary Fig. 1) confirms the reciprocal monophyly of seven distinct subspecies clusters with Bootstrap support of e 80% (Bock et al., 2014; Brown et al., 2007; Fennessy et al., 2016). MtDNA do not support two subspecies of Masai giraffe because individuals which are designated Masai giraffe individuals, disrupt a possible reciprocal monophyly. For reticulated giraffe, three individuals do not group as expected but rather fall within the northern giraffe, indicating possible hybridization. However, two of these individuals are from zoos, where hybridization is common, and have an unknown history. The third individual is a wild giraffe from a geographic range adjacent to the northern giraffe and is a possible natural hybrid, but during sampling was identified as a phenotypically reticulated giraffe.

**Fig. 2.**
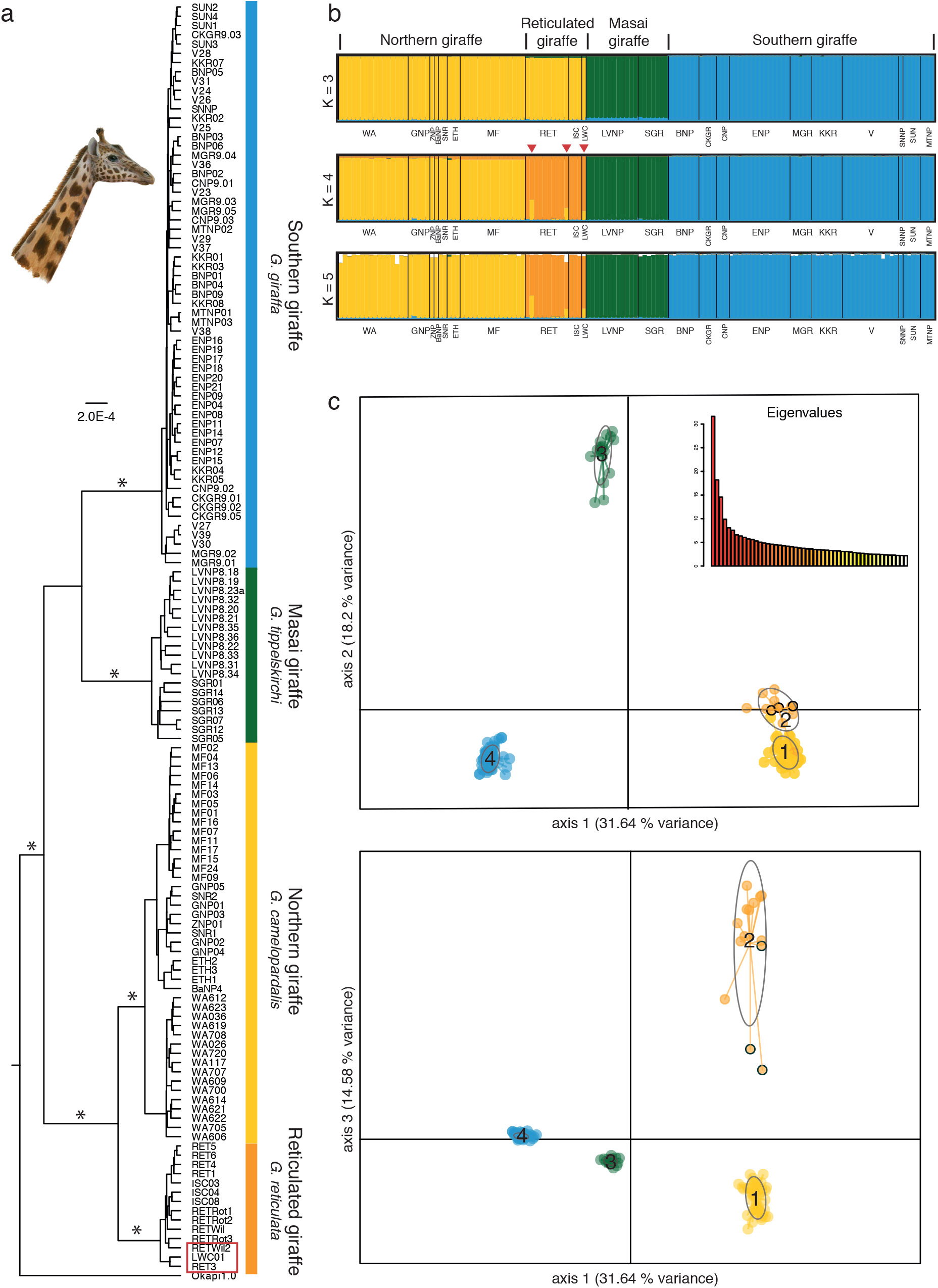
Nuclear phylogeny and population structuring of giraffe. (a) Bayesian multi-locus tree from 21 nuclear loci and 137 giraffe individuals reconstruct four significant supported (p ≥ 0.95) giraffe clades, corresponding to the four giraffe species (Fennessy et al., 2016). The okapi is used as the outgroup. The asterisks indicate branches with statistical significant support (p ≥ 0.95). The red frame indicates the potential hybrids. (b) STRUCTURE analysis of the dataset, excluding the okapi. The colors indicate the membership in a cluster for each sampling location and individual. K = 4 shows four well-resolved groups and is supported as best fitting number of clusters by several statistical methods (see Supplementary Fig. 2). The grouping into four clusters is consistent with the Bayesian multi-locus analysis: yellow: northern giraffe, orange: reticulated giraffe, green: Masai giraffe, and blue: southern giraffe. Three individuals within the reticulated giraffe cluster (red arrowheads) indicate potential hybridization with admixture from the northern giraffe. K = 3 merges northern and reticulated giraffe, and at K ≥ 5 no further clustering is evident. (c) PCA axes 1-2 and axes 1-3 for four distinct giraffe clusters (1: northern; 2: reticulated; 3: Masai; 4: southern). Colors as in Fig 2b. The 95% confidence intervals are shown as oval outlines. Note that the non-overlapping confidential intervals in the PCA axes 1-2, as well as, axes 1-3 indicate significantly different clusters. Potential hybrids are indicated by black circles. Note – The drawing by Jon B. Hlidberg shows a Nubian giraffe.

Multi-locus population STRUCTURE analyses (Pritchard, Stephens, & Donnelly, 2000) of 21 nuclear loci (Fig. 2b) proposes the best clustering into four distinct populations (optimal K = 4) based on the graphical display. At K = 3 the analyses merge the reticulated and the northern giraffe and at K ≥ 5 the analyses do not produce further clustering. Three different statistical methods to interpret the STRUCTURE results (Evanno, Regnaut, & Goudet, 2005; Pritchard et al., 2000; Pritchard, Wen, & Falush, 2010) confirm K = 4 being significantly the best fitting number of populations (Supplementary Fig. 2). These four clusters conform with the four giraffe clades identified by tree analyses. Intriguingly, STRUCTURE also identifies three potential hybrids between the northern and reticulated giraffe within the reticulated giraffe clade (Fig 2a, Fig 2b). The distinctness of four unique giraffe clades is in addition supported by Principal Component Analyses (PCAs) (Fig. 2c) with significant non-overlapping 95% confidential intervals. PCAs using groups of the seven mtDNA clades do not find more than four distinct clusters (Supplementary Fig. 3). Finally, pairwise fixation indices (F_st_) of ≥ 0.237 (statistically significant at p < 0.001) are consistent with the four distinct clusters of giraffe in the tree analyses (Supplementary Table 3).

Separate PCAs and STRUCTURE analyses for each species (Supplementary Fig. 4-7) indicate population substructure within northern giraffe, and potentially in the Masai giraffe, but no further population substructure in southern and reticulated giraffe. Within northern giraffe STRUCTURE and PCAs up to four clusters can be identified. However, the sample sizes of some populations (three for Ethiopia) are arguably insufficient to draw definitive conclusions. Within the Masai giraffe STRUCTURE and PCAs identify potentially two separate clusters, indicating a possible separation of the two geographically most distant populations that have been analyzed for nuclear SNPs (single nucleotide polymorphisms) to date. Consistent with the STRUCTURE and PCA analyses, pairwise F_st_ analyses within each giraffe species finds a high level of population differentiation within northern and possibly Masai giraffe, and little differentiation within southern and reticulated giraffe (Supplementary Table 4).

We estimated long-term gene flow within all four giraffe clades, as well as among subspecies within each of the giraffe species which show population substructure in STRUCTURE and PCAs using MIGRATE-N (Beerli, 2006; Beerli & Felsenstein, 2001). Assuming similar mutation rates among all giraffe species, the mutation-scaled population size theta (Θ) estimates for the four species suggest that the effective population size (N_e_) is smaller in southern giraffe and Masai giraffe than in northern and reticulated giraffe (Supplementary Table 5a). Thus, the population size in the northern and reticulated giraffe had been larger in the past. The calculated effective numbers of migrants per generation or gene flow rate (N_e_m) based on Θ and the mutation-scaled migration rate (M) (Supplementary Table 5b) indicate generally very low level of gene flow among most of the four giraffe clades with a maximum of one migrant per five generations (N_e_m < 0.179), with one exception. A higher N_e_m occurs between the northern and reticulated giraffe, with nearly one migrant per generation in the direction of the reticulated giraffe (N_e_m = 0.945), but much less migration is observed in the opposite direction to northern giraffe (N_e_m = 0.179). There is little (ca. one in ten) directional gene flow from Masai to reticulated giraffe (N_e_m = 0.107) and from southern to reticulated giraffe (N_e_m = 0.104) with nearly zero gene flow in the opposite direction. The gene flow rates for all other species pairs are extremely low (N_e_m < 0.065). Within species long-term gene flow rates are on average higher (N_e_m >1) (Supplementary Table 6b). However, between some subspecies gene flow is also limited, in particular the geographically extremely isolated West African giraffe (WA).

Gene flow rates that were calculated on pairwise F_st_ values between species corroborate the MIGRATE-N analyses, but provide no information about the direction of gene flow and the rates are somewhat higher (Supplementary Table 3). In agreement with the MIGRATE-N analyses, the F_st_ based analyses find the highest rate of gene flow between northern and reticulated giraffe (N_e_m = 0.804), and a much lower gene flow rate (N_e_m ranges from 0.113 to 0.186) is observed among all other population pairs.

Finally, short-term migration rates (m) estimated with BayesAss (Wilson & Rannala, 2003) (Figure 3, Supplementary Table 5b) confirm low levels of gene flow among the four giraffe species for the past three generations. The highest migration rates occur from northern, Masai and southern giraffe in the direction of reticulated giraffe. The data suggest that approximately 2% (m = 0.021) of the reticulated giraffe population are derived from each of these neighboring species, which is expected. In comparison, to the other gene flow analyses, BayesAss identifies somewhat higher recent migration rates (m) among subspecies within species (Supplementary Table 6b). This is consistent with the lack of genetic differentiation identified by PCA and F_st_ analyses. The recent migration rates estimated by BayesAss analyses suggest directional gene flow between West African and Kordofan giraffe (m = 0.064), and find gene flow between South African and Angolan giraffe (m = 0.052). Most importantly, however is that BayesAss does not find any first or second generation hybrids.

**Fig. 3.**
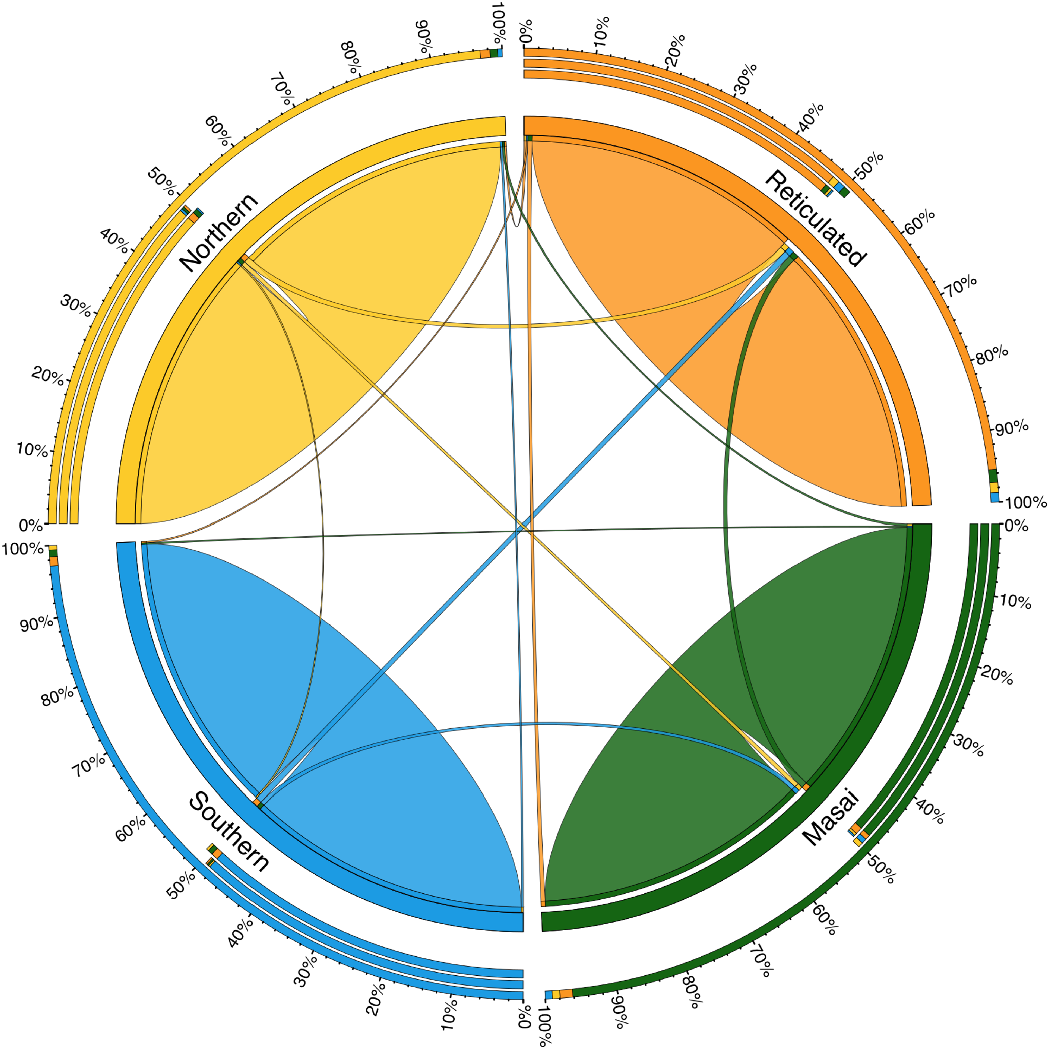
Circular migration plot of recent migration rates among four giraffe clades. Recent directional migration rates (m) as estimated by BayesAss and indicated by ribbons connecting one species to another. The color coding of the four species is according to the STRUCTURE clusters (Fig 2b). Peripheral concentric stack bars show relative migration rates in percent. Whereas the inner stack bar shows the outgoing ribbon sizes, the middle stack bar the incoming ribbon sizes and the outer stack bar the combination of both.

## Discussion

Morphology and genetic analyses suggest that there is more than one giraffe species (Brown et al., 2007; Fennessy et al., 2016; Groves & Grubb, 2011). Here we expand our previous dataset three-fold and improve the sampling of northern and reticulated giraffe, to further study if there are indeed more than one species. The new data set allows for the first-time detailed gene flow and migration analyses. Among the four giraffe species^6^, gene flow and migration is very limited. As such, the new analyses of the extended nuclear data corroborate the identification of four genetically distinct giraffe species (Fennessy et al., 2016).

Several attempts have been made to define a species, but a unequivocal consensus has not yet been reached (Coyne & Orr, 2004; De Queiroz, 2007). The most commonly applied model is the BSC, which suggests that reproductive isolation is essential to delineate species (Dobzhansky, 1970; Mayr, 1942). By contrast, subspecies or evolutionary significant units (ESU) are sometimes arbitrary distinctions within a species. Reproductive isolation is also a cornerstone of other species concepts that define species as distinct ESUs with limited gene flow to other such units (Avise & Ball, 1990). Therefore, analyzing gene flow among species is a central analysis to delineate species, especially if, like in giraffe, they possibly hybridize in nature. It has been suggested that gene flow among species must be limited to allow genetic differentiation, and a value below one migrant per generation (< 1 N_e_m) is a conservative estimate (Wright, 1969), even if other studies are more liberal and suggest that gene flow rates of < 5 N_e_m (Lacy, 1987) or even < 10 N_e_m (Mills & Allendorf, 1996; Vucetich & Waite, 2000) can allow genetic differentiation and consequently speciation.

The initial finding of four giraffe species was unexpected (Fennessy et al., 2016), because giraffe seem to be a morphologically homogenous group, can interbreed in captivity (Gray, 1972; Lackey, 2011; Lönnig, 2011), and are highly mobile (Flanagan, Brown, Fennessy, & Bolger, 2016). To avoid differentiation between populations, and if giraffe was in fact one species, gene flow rates in excess of 1-10 migrants per generation are expected based on mathematical models (Lacy, 1987; Mills & Allendorf, 1996; Vucetich & Waite, 2000; Wright, 1969).

Our study show that the long-term average estimates of gene flow rate among giraffe are below one migrant per generation (N_e_m < 1). The introgression and population genetic analyses of the expanded data set are thus consistent with the previously proposed classification of four giraffe species by Fennessy et al. (2016). The highest gene flow rates are observed between the northern and reticulated giraffe, but the rate is below Wright’s (1969) conservative estimate of < 1 N_e_m. Compared to other giraffe species, higher gene flow rates among northern and reticulated giraffe is not unexpected, because they are the closest related (Fennessy et al., 2016) and their current neighboring geographic ranges might have overlapped historically (Fig. 1).

Yet, even among the closely related and neighboring northern and reticulated giraffe lower gene flow rates than < 1 N_e_m are observed, which is consistent with being genetically differentiated species. In addition, among all 137 individuals from a wide geographic distribution, only one natural hybrid has been genetically identified yet. The rare occurrence of hybrids further supports the existence of four giraffe species.

Population genetic analyses, such as STRUCTURE and PCA of the data set support the results from the gene flow analyses. The new results are inconsistent with past suggestions of possibly six or seven distinct giraffe species (Brown et al., 2007). These results were based on non-stringent conclusions from STRUCTURE analyses with 11 separate genetic clusters at K = 13 based on 14 microsatellites, and results of six to seven giraffe clusters based on mtDNA phylogeny (Brown et al., 2007): “11 of the 18 sampling localities resolved as distinct genetic clusters at K=13”, however the authors concluded that only “the seven lineages that are reciprocally monophyletic in the mtDNA tree need to be considered evolutionary significant units if not species”. Other findings of up to eight giraffe species were proposed based on a combination of limited genetic analyses (Brown et al., 2007; Hassanin, Ropiquet, Gourmand, Chardonnet, & Rigoulet, 2007) and morphological characteristics (Groves & Grubb, 2011), however the location of some samples were inaccurate.

Both Fennessy et al. (2016) and Bock et al. (2014) suggested to subsume Rothschild’s giraffe (MF) into the Nubian giraffe, as well as Thornicroft’s giraffe (LVNP) into the Masai giraffe, because they lack differentiation at mtDNA sequences. Evolutionary differentiation of populations is often first evident in mtDNA, because theory suggests that this locus, due to its maternal inheritance and non-recombining nature, reaches fixation 4-times more rapidly than nuclear loci (Zink & Barrowclough, 2008). Thus, differentiation into subspecies is often first evident on mtDNA, rather than nuclear sequences. Such population differentiation processes have been reported in natural population of bears (Hailer et al., 2012), humpback whales (Palumbi & Baker, 1994) and macaques (Melnick & Hoelzer, 1992).

While the current mtDNA analyses support previous findings (Fennessy et al., 2016) of Thornicroft’s giraffe being subsumed into the Masai giraffe, new and extended nuclear gene datasets identify some substructure among them. We emphasize however, that the nuclear loci have only been sampled from across a limited distribution of the Masai giraffe^6^. Additional sampling of intermediate Masai giraffe populations and additional nuclear gene loci will be necessary to yield more definite results. The first detailed mtDNA analyses on Thornicroft’s giraffe (Fennessy, Bock, Tutchings, Brenneman, & Janke, 2013) proposed that while they are not reciprocal monophyletic, the geographic location in Zambia’s Luangwa Valley is unique and should, for conservation efforts, tentatively maintain its subspecies status as Thornicroft’s giraffe within Masai giraffe (*Giraffa tippelskirchi thornicrofti*).

Within the northern giraffe some substructure is evident in PCAs and STRUCUTURE analyses for nuclear sequences (Supplementary Fig. 6). However, the West African giraffe is a geographically very isolated and small population of ∼600 individuals. As described (Fennessy et al., 2016), the geographic distinction between the former Nubian and Kordofan giraffe is unclear and current data suggest that they are not genetically isolated.

Multi locus phylogenies, population genetic and gene flow analyses support the hypothesis of four genetically distinct giraffe species (Fennessy et al., 2016). The molecular data show that there is only very limited gene flow between the four species, which is in agreement with the BSC.

With little more than 5,000 northern giraffe, <8,700 reticulated giraffe and ∼34,000 Masai giraffe remaining in the wild (Giraffe Conservation Foundation, 2017), recognizing these – and the southern giraffe – as separate species has an impact on giraffe conservation. Their decline in numbers over the last thirty years (three generations) – northern giraffe (∼95 %), reticulated giraffe (∼80%) and Masai giraffe (∼52 %), highlight that these species are threatened with extinction(IUCN, 2017). Giraffe, as a single species, and not four, were recently listed as “Vulnerable” on the IUCN Red List (Muller et al., 2016). The mounting evidence of four giraffe species proposes a re-evaluation of the current IUCN giraffe taxonomy to raise the conservation classification to a higher level of threat, and in turn increased conservation management actions.

## Acknowledgements

We thank Susanne Gallus, for support in the laboratory, and Fritjof Lammers as well as Markus Pfenninger for support with analyses. Stephanie Fennessy and Maria Nilsson gave valuable comments on the manuscript. We thank Jon B. Hlidberg for the artwork of a Nubian giraffe. This study was supported by the Leibniz Association, the Giraffe Conservation Foundation-USA, the Leiden Conservation Foundation, and various African government partners and international zoo and private supporters.

## Data Accessibility Statement

The authors declare that all the data supporting the findings of this study are available within the article and its Supplementary Information files. Sequences generated during the study are available at GenBank (https://www.ncbi.nlm.nih.gov/nucleotide/) under the Accession Numbers MG257948 – MG262301.

## Author Contributions

AJ, JF and SW designed and conceived the study. AJ and JF funded the project, JF collected the samples and provided biological data. SW developed markers, generated and analyzed the data. SW and AJ wrote the manuscript with input from JF.

